# A methodological approach to developing evidence-based communication guidelines for *Populus* spp. cultivation on agricultural land

**DOI:** 10.1101/2025.08.11.669630

**Authors:** Kristina Blennow, Elin Anander, Johannes Persson, Annika Wallin

## Abstract

This study developed and applied a methodology and framework to analyse decision-making and identify communication needs related to *Populus* spp. cultivation on agricultural land in southern Sweden, where adoption remains limited despite significant biomass potential. By comparing farmers’ willingness and actual cultivation decisions, and linking these to expectations on economic, agronomic, and environmental outcomes, we moved beyond attitudes to analyse choices. We used Bayesian Additive Regression Trees (BART) to predict cultivation decisions and willingness to cultivate using Net Value of Expected Impact (NVEI) as a predictor. Integrating decision-making with willingness revealed drivers of adoption and barriers persisting despite positive attitudes. NVEI scores strongly predicted both willingness and cultivation decisions, with regional contrasts evident. In the Forested region, farmers anticipated greater economic and agronomic benefits but had lower expectations for biodiversity, suggesting a knowledge gap and a need for communication providing comparative evidence on biodiversity benefits. In the Lowland region, higher expectations for environmental benefits highlight the need for tailored information on biodiversity, landscape value, and ecological connectivity. Low scores for workload and management ease across all groups indicate uncertainty about practical aspects, supporting the need for targeted communication. Overall, regionally tailored, evidence-based guidelines can support informed decision-making.

## 1. Introduction

In the pursuit of climate neutrality by 2050, the European Union (EU) has adopted targets under the European Green Deal (European Council 2025) and the revised Renewable Energy Directive (RED III), calling for at least 42.5% of the EU’s gross final energy consumption to come from renewable energy sources by 2030 (EU 2023). Biomass, the EU’s largest source of renewable energy (EEA 2025), is expected to play a central role in this transition, not only in replacing fossil fuels for energy but also as a key feedstock for bio-based materials and products. However, the potential to expand biomass production sustainably hinges on aligning technical, environmental, and economic assessments with the practical realities and decisions of landowners, particularly farmers.

Wood from short-rotation tree species such as *Populus* spp. is highlighted in policy frameworks as a strategic feedstock due to its high productivity and potential to avoid conflicts with food production (see e.g. Naess et al. 2023). Yet, despite technical assessments that identify large biophysical potentials for this resource, actual market availability remains constrained. Prior studies have demonstrated that land often classified as “available” in top-down models, such as abandoned or underutilised agricultural land, may still serve important functions for farmers, or may not be perceived as viable for biomass production (e.g. Anander et al. 2024; Thomson Ek et al. 2024).

The underrepresentation of farmers’ perspectives in biomass potential assessments entails a risk of misrepresentation of biomass supply, leading to misinformed policy expectations (e.g. Blennow et al. 2014; Anander et al. 2024; Blennow et al. submitted). Several studies highlight the role of market uncertainties and price sensitivity in shaping farmers’ willingness to produce or sell biomass, particularly for *Populus* spp. and other short-rotation woody crops (Schulze et al. 2016, Thomson Ek 2024). Paulrud and Laitila (2010) found that farmers’ attitudes, rather than purely economic benefits, played a significant role in the willingness to cultivate energy crops on agricultural land. For example, in a choice experiment, the production of taller crops and longer rotation periods, such as with *Populus* spp. compared to annual crops, significantly decreased the farmers’ expected utility of the crop. Similarly, a sceptical and negative attitude toward the cultivation of trees on agricultural land for bioenergy among U.S. Great Plains farmers has been reported (Hand and Tyndall 2018). In addition, Wilson et al. (2014) found that English farmers’ reluctance to consider growing dedicated energy crops was often due to concerns about committing land over a long time period. Nevertheless, current bioenergy strategies rarely consider farmers’ informational needs or decision-making contexts, as seen, for example, in Böhlenius et al. (2023), factors that are essential for fostering participation in biomass markets and ensuring realistic supply projections.

To support the transition to a sustainable bioeconomy, it is essential to bridge the communication gap between biomass policy ambitions and farmers’ informational needs. While earlier studies have explored economic and environmental dimensions of perennial energy crops, relatively little is known about how farmers’ prior experience and contextual factors, such as regional differences, shape both their willingness and actual decisions to cultivate *Populus* spp. on agricultural land. Understanding these drivers is key to aligning policy and communication strategies with on-the-ground reality.

This study aims to develop and apply a methodology for identifying evidence-based communication needs related to the cultivation of *Populus* spp. on agricultural land. By empirically investigating how prior experience with silviculture relates to both willingness and decisions to cultivate *Populus* spp., we illustrate how a decision-making (analytic) framework can inform targeted communication strategies and policy design. The empirical case focuses on southern Sweden, where, despite high theoretical potential for biomass supply from *Populus* spp., adoption rates remain low, although regional variation in attitudes and adoption exists.

Using a *problem-feeding* approach (Thorén and Persson 2013; Persson et al. 2018), we developed and applied a methodology and framework to analyse decision-making to inform evidence-based communication about biomass crop adoption. Drawing on earlier work (Anander et al. 2024), we first used interviews to identify key challenges as perceived by farmers in practical agriculture, particularly regarding the cultivation of *Populus* spp. We then assessed the relevance and extent of these challenges through a regional survey.

The survey results revealed that farmers in southern Sweden generally held less favourable views on cultivating *Populus* spp. than previously estimated in biomass potential studies (Anander et al. 2024). However, the data also showed clear regional variation: willingness to cultivate hybrid aspen or poplar was significantly higher among respondents in forested regions compared to those in intermediate or lowland areas.

To interpret and structure these findings, we applied a survey-based framework for analysing decision-making, developed in previous studies on climate change decision-making (Blennow et al. 2020, 2021; Persson et al. 2020; Blennow and Persson 2021), that estimates the decision-making agents’ expected utility (*sensu* von Neumann and Morgenstern 1944). By analysing both actual and potential decision-making, we demonstrate how this methodological framework can be used to identify critical determinants of farmers’ willingness and behaviour, and how it supports the development of targeted communication strategies.

We test the following hypotheses:

Net expected economic, agronomic, and environmental impacts and consequences of cultivating *Populus* spp. are correlated with actual decisions to cultivate or expand the cultivation of *Populus* spp. on agricultural land.

1. Net expected economic, agronomic, and environmental impacts and consequences of cultivating *Populus* spp. are correlated with farmers’ willingness to cultivate *Populus* spp. on agricultural land.
2. Previous experience with silviculture or tree cultivation is positively correlated with actual decisions to cultivate or expand the cultivation of *Populus* spp. on agricultural land.
3. Previous experience with silviculture or tree cultivation is positively correlated with farmers’
4. willingness to cultivate *Populus* spp. on agricultural land.

Hypotheses 3 and 4 are interesting to consider in light of both the description/experience-distinction (Hertwig et al.2018), which is often applied to learning, decision-making and adaptation, and the notion of science and proven experience which has had a formative role in Swedish thinking about the intersection between academic and experiential knowledge (e.g. Persson et al. 2019).

Empirical consequences of these hypotheses were tested relating to expected utility components classified as economic (expected consequences on *Farm profitability, Property market value, Alternative land use*, and *Risk spreading*), agronomic (expected impacts on *Farm workload, Long rotation, Easy to manage, Resprouting ability*, and *Suitability in a changing climate*), or environmental (expected impacts on *Biodiversity* and *Landscape character*) and indicators of experience of silviculture and tree cultivation.

## Materials and methods

### 2.1 Study area

Scania County, located in southern Sweden (Figure 1), is characterised by its extensive agricultural land coverage, about 45% of the total area (Statistics Sweden 2023), and a high density of agricultural enterprises. The county’s arable land is the most fertile in Sweden, with agricultural production largely centred on annual crops, particularly cereals (Swedish Board of Agriculture 2025). However, land use varies significantly across the region. In the lowland coastal areas, agriculture dominates the landscape, covering 60–81% of the land, whereas the northern central areas are predominantly forested, with only 5–19% agricultural land. The intermediate zone features a more balanced mix of forest and farmland (Swedish Board of Agriculture 2025) (Figure 1). This variation in land use is closely linked to differences in soil productivity: the arable lowlands are characterised by fertile, high-yielding soils, while soils in the forested uplands and transitional areas are generally less productive and less suitable for intensive crop cultivation (Erlandsson 1999). The value of land is higher in the Lowland and Intermediate regions than in the Forested region (Swedish Board of Agriculture 2019).

**Figure 1.**
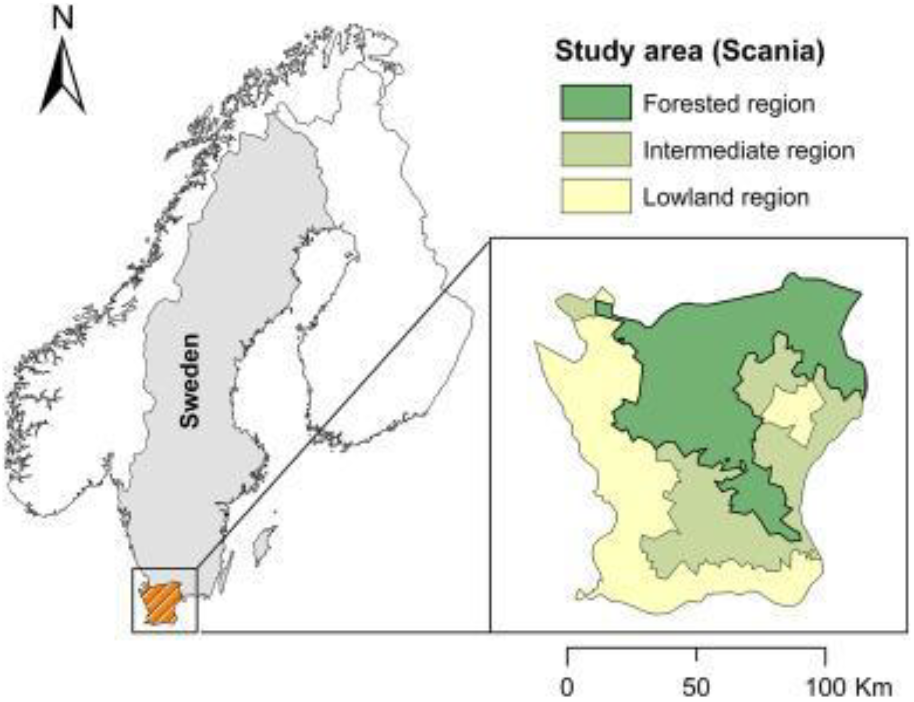
The study area, Scania, is located in southern Sweden. The colour coding indicates three dominant types of land use. Made with data from Natural Earth and reproduced from Anander et al. (2024).

Animal husbandry, especially cattle and milk production, is most common in the forested and intermediate zones, and least prevalent in the lowland areas (Erlandsson 1999). In 2020, Scania’s agricultural land totalled 490,000 ha, of which arable land, excluding pasture land, totalled 435,000 hectares, representing 17% of Sweden’s total arable area of 2.5 million hectares (Swedish Board of Agriculture 2025). In the same year, 798 hectares of agricultural land in Scania were cultivated with hybrid aspen and poplar, corresponding to 0.16% of the county’s total agricultural land or 0.18% of its arable land (Swedish Board of Agriculture 2025).

### 2.2 Data collection

#### 2.2.1 Survey design and sample selection

Building on insights obtained from an explorative pre-study, a broad, unstratified internet-based survey was conducted to obtain diverse samples of farmers concerning regional location (Anander et al. 2024). Approximately 50% of members of the Federation of Swedish Farmers (LRF) in Scania were randomly sampled and invited via e-mail by LRF to participate in the survey (n=2,567). The survey addressed short-rotation *Populus* spp. cultivation. *Populus* spp. includes, among others, hybrid aspen and various poplar species. The parent LRF membership group represents approximately 85% of all agricultural holdings registered in Scania’s farm register in 2020 (Swedish Board of Agriculture 2025).

The survey was conducted in Swedish, using the Netigate survey tool (2024) for both development and distribution. The questionnaire was accompanied by a cover letter explaining the study objectives and emphasising that participation was voluntary (see Anander et al. 2024).

#### 2.2.2. Data collection and response rate

The survey was deployed on February 17, 2021 and remained open until April 23, 2021. A reminder about participation was issued once to all farmers in the sample.

The survey, which comprised 30 questions, is described in detail in Anander et al. (2024). Responses to seven of the 30 questions were used for data analysis in this study (Table 2).

A total of 179 respondents submitted responses to the questionnaire, yielding a response rate of 7%. While this may appear modest, it is consistent with similar survey-based research involving farmers and landowners across Europe (e.g., Rommel et al. 2022). Such studies frequently report low participation rates due to the long-term nature of the decision, limited familiarity with the subject, or time constraints among the farming population. Low response rates are therefore typical in this research area and not necessarily indicative of low data reliability.

Of the 179 responses in this study, the responses of 73 respondents were excluded due to incomplete information. A refined sample of 106 responded to the questions used in this study (4% response rate), representing 2% of the farming community in Scania. The questions analysed for this study focus on farm location and respondents’ values of expected impacts of cultivating *Populus* spp. for energy use (Table 1). Responses to Q.3 in Table 1 was also analysed in Anander et al. (2024).

**Table 1.** Questions analysed in the present study on supplying biomass from *Populus* spp. See Anander et al. (2024) for the complete questionnaire.

| Number | Question | Response option |
| --- | --- | --- |
| Q.1 | In which harvest zone is the land you cultivate located? <sup>a</sup> | Aggregated into three regions:<br>Forested region<br>Intermediate region<br>Lowland region<br>Do not know |
| Q.2 | Have you been involved in the decision to plant any of these areas<sup>b</sup> with hybrid aspen/poplar? | Yes<br>No |
| Q.3 | During the upcoming 10-year period 2021–2030, do you think you will plant/plant more stands of hybrid aspen or poplar on your farm's agricultural land?<br>a. Hybrid aspen<br>b. Poplar | No, definitely not<br>Probably not<br>Don't know<br>Yes, probably<br>Yes, definitely |
| Q.4 | On your farm's agricultural land, do you expect the cultivation of hybrid aspen or poplar to:<br>benefit farm profitability<br>increase the value of the property<br>provide alternative land use<br>help spread risk<br>reduce farm workload<br>benefit your enterprise because of its 20-year rotation period<br>be advantageous due to ease of management<br>be beneficial because of its ability to resprout from the root or stump<br>be beneficial because of its suitability to a changing climate | Yes, always<br>Often<br>Seldom<br>No, never<br>Don't know |
|  | • benefit biodiversity<br>• enhance the landscape character |  |
| Q.5 | On your farm's agricultural land, do you expect the cultivation of hybrid aspen or poplar to:<br>• not benefit farm profitability<br>• not increase the value of the property<br>• not provide an alternative land use<br>• not help spread risk<br>• not reduce farm workload<br>• not benefit your enterprise because of its 20-year rotation period<br>• not be advantageous due to ease of management<br>• not be beneficial because of its ability to resprout from the root or stump<br>• not be beneficial because of its suitability to a changing climate<br>• not benefit biodiversity<br>• not enhance the landscape character | Yes, always<br>Often<br>Seldom<br>No. never<br>Don't know |
| Q.6 | On your farm<sup>c</sup>: Is there currently any land planted with stands of hybrid aspen or poplar? | Yes<br>No<br>Don't know |
| Q.7 | Do you own any forest land? | Yes<br>No |

<sup>a</sup> The detailed farm locations were initially collected based on multiple harvest zones. For this analysis, these zones were aggregated into three broader regions: Forested region, Intermediate region, and Lowland region (see Anander et al. 2024). <sup>b</sup> Any of these areas" refers to the question "What is the age of your stand(s) of hybrid aspen/poplar?" not analysed in this study.
<sup>c</sup> Cultivation of *Populus* spp. on forest land was excluded from the present study. The exclusion was based on a follow-up question on the type of land used for cultivation.

**Table 2.** Strength of evidence based on posterior probability (Jeffreys 1939).

| Evidence strength | Posterior probability |
| --- | --- |
| Null hypothesis supported | $< 0.5$ |
| Bare mention | $0.50 \leq p < 0.77$ |
| Substantial evidence | $0.77 \leq p < 0.91$ |
| Strong evidence | $0.91 \leq p < 0.97$ |
| Very strong evidence | $0.97 \leq p < 0.99$ |
| Decisive evidence | $\geq 0.99$ |

### 2.3. Data preparation

Following Persson et al. (2020) and Blennow et al. (2020), we calculated the *Net Value of Expected Impacts* (NVEI) for the decision and willingness to cultivate *Populus* spp. for energy use (Q.4 and Q.5 in Table 1). NVEI was calculated by summing the net scores of paired positive and negative expectations regarding biomass provision decision and willingness (Q.2 and Q.3 in Table 1, respectively).

For each expectation, responses of “Yes, always” were assigned values of +4 (positive) or −4 (negative), “Often” responses were assigned +3 or −3, respectively, and all other response options (“Seldom”, “Never”, “Don’t know”) were assigned 0. These values were summed across all items for each respondent to produce the NVEI score. We also computed *partial NVEI* scores, which represent the contribution of individual sub-questions (i.e., specific expectations).

### 2.4 Statistical analysis and machine learning modelling

Bayesian methods were used due to their robustness with small and unevenly distributed samples. Unlike frequentist approaches, Bayesian inference can yield credible parameter estimates and posterior probabilities without requiring large sample sizes (McElreath 2016). It also allows for direct probabilistic statements about group differences and model comparisons. All analyses were conducted in R version 4.4.1 (R Core Team, 2024).

We used the interpretation scale by Jeffreys (1939) for evaluating the strength of evidence (see Table 2). Weakly informative priors were applied to stabilise estimation while maintaining minimal influence on posterior distributions, preserving interpretability under Jeffreys’ scale.

Following the Bayesian framework and drawing on Jeffreys’ (1939) interpretation of evidence strength, we define strong support for a hypothesis as a posterior probability equal to or greater than 0.91.

#### 2.4.1 Machine learning modelling

We applied Bayesian Additive Regression Trees (BART) to model the probability of cultivation and willingness to cultivate *Populus* spp., using NVEI scores as predictors, with the *bartMachine* package in R (Kapelner and Bleich, 2016). BART is a non-parametric machine learning method that builds an ensemble of shallow regression trees, where each tree contributes incrementally to the prediction. It uses a Bayesian framework to regularise the model and prevent overfitting. To evaluate model performance and predictive accuracy, we implemented 3-fold cross-validation, repeating the procedure three times with different random splits to ensure stability of results. In this study, BART was used for classification, and its probabilistic predictions were interpreted as the likelihood of making, or being willing to make, the biomass provision decision, respectively.

Model performance was evaluated using the Area Under the Curve (AUC) of the Precision-Recall (PR) curves, representing the model’s ability to correctly identify positive cases across various thresholds while accounting for class imbalance. PR curves are particularly informative in this context, as they focus on the minority (positive) class and provide a more realistic assessment than ROC curves when positive cases are rare. The PR curves and AUC calculations were performed using the PRROC package in R (Grau et al., 2015). Model interpretation was supported by Partial Dependence Plots (PDPs) generated using the ICEbox package (Goldstein et al., 2015).

#### 2.4.2 Hypothesis testing

To test for differences between groups, such as regions, we used the Bayesian Estimation Supersedes the t-Test (BEST) approach. Models were implemented using the BEST package (Kruschke 2015) in R.

Differences in proportions between groups were tested using a Bayesian proportion test, implemented with the Bayesian First Aid package in R (Bååth 2014).

## 3. Results

### 3.1. Predictive modelling of the decision and willingness to cultivate *Populus* spp. on agricultural land

Of 106 respondents, only five reported having participated in a decision to cultivate *Populus* spp. on agricultural land (Q.2 in Table 1). A univariate BART model showed that the probability of such a decision increases sharply when the NVEI exceeds a threshold of 30 (Figure 2a). The model demonstrated excellent classification performance, with an area under the precision-recall curve (PRAUC) of 0.87 (Figure 3a).

**Figure 2.**
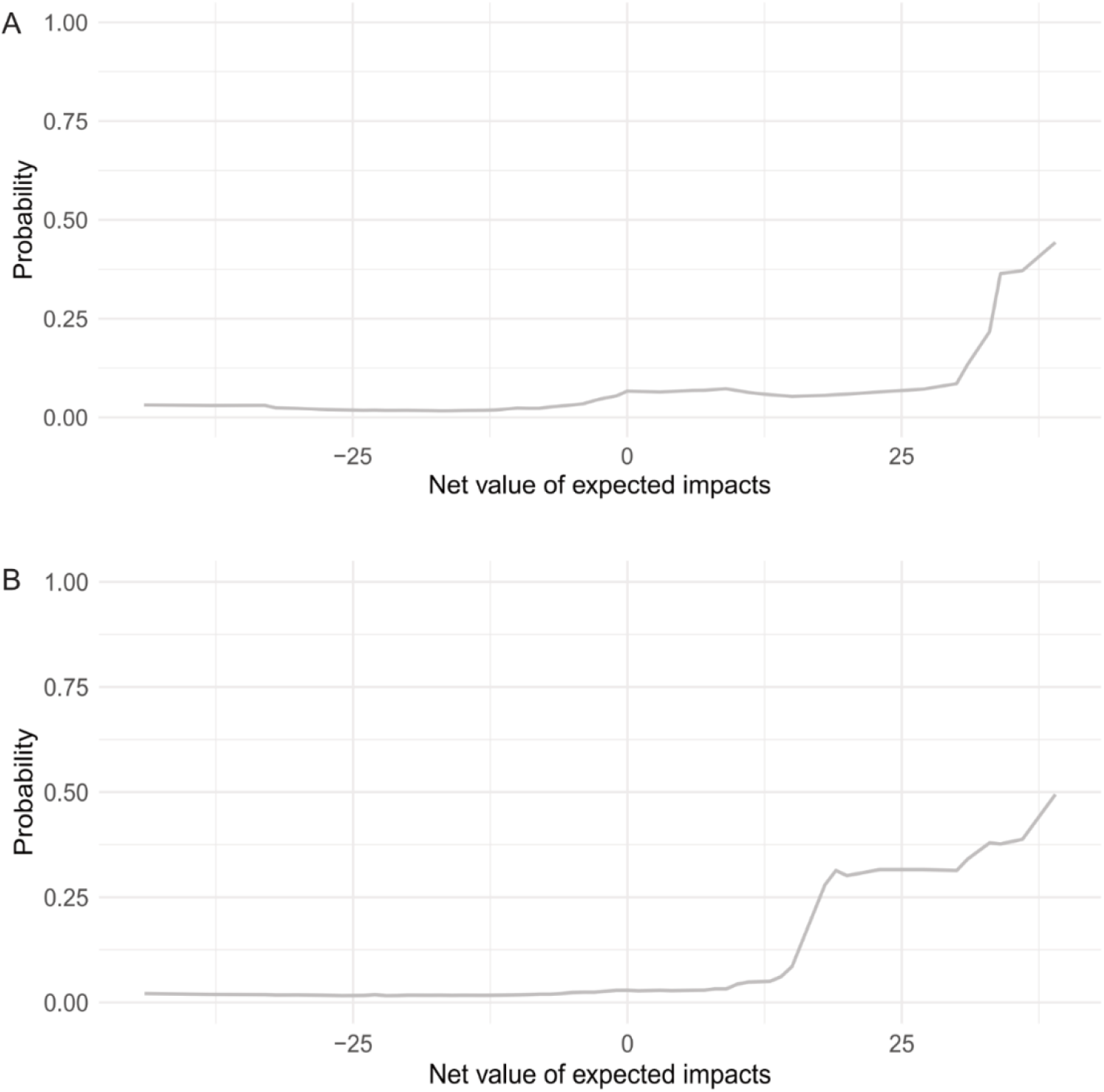
Partial Dependence Curve (PDP) of the relationship between NVEI and the predicted probability of (a) having decided to cultivate *Populus* spp. on agricultural land (Q.2 in Table 1) and (b) being willing to consider cultivating *Populus* spp. on agricultural land in the decade 2021–2030 (Q.3 in Table 1), based on univariate BART models.

**Figure 3.**
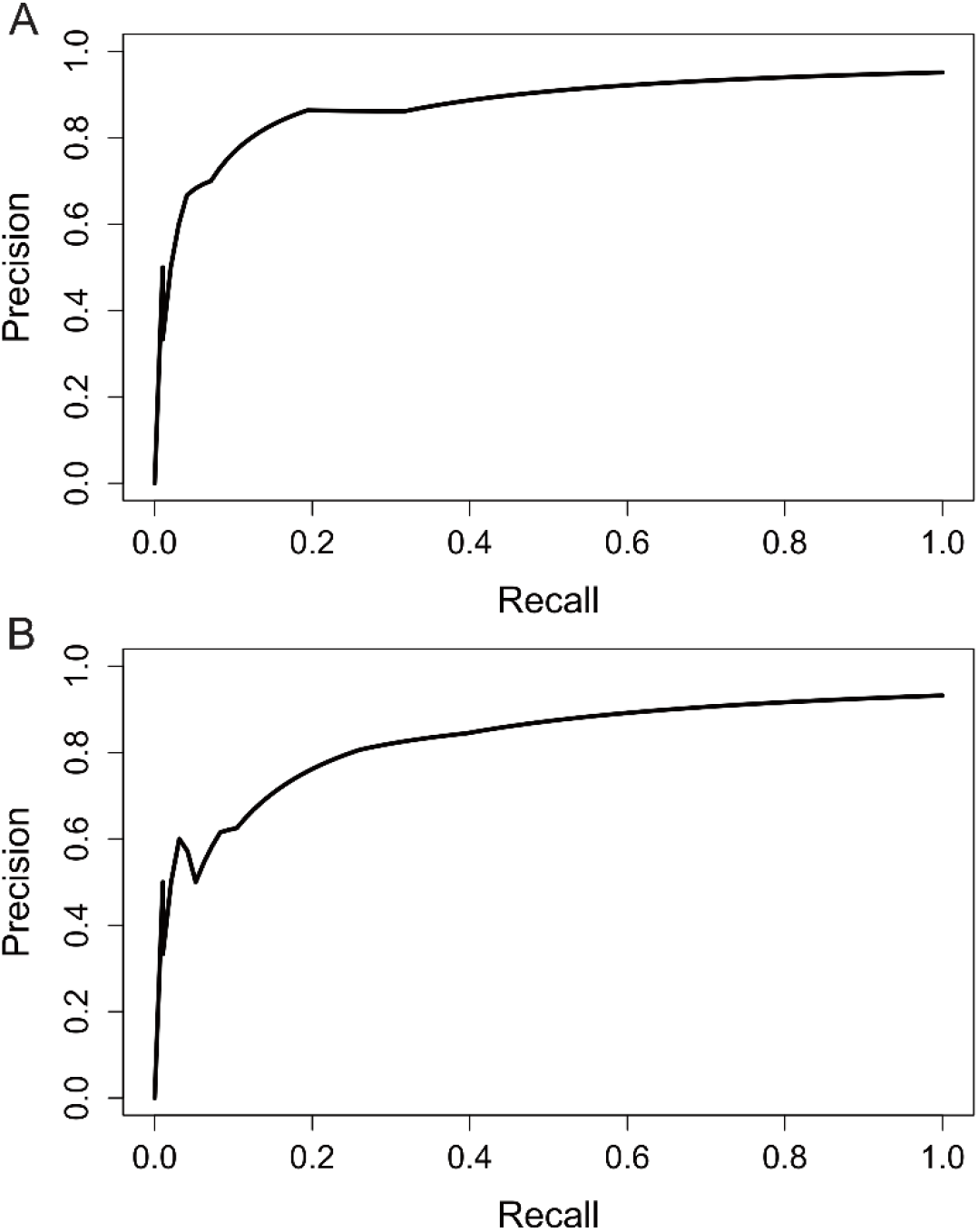
Precision-recall (PR) curves for the univariate BART models: (A) predicting whether respondents had decided to cultivate *Populus* spp. on agricultural land (Q.2 in Table 1), with a PR area under curve (PRAUC) of 0.87; and (B) predicting willingness to consider cultivating or expanding cultivation of *Populus* spp. on agricultural land within the coming decade (Q.3 in Table 1), with an PRAUC of 0.83.

For willingness, ten respondents indicated they would consider initiating or expanding *Populus* spp. cultivation within the next decade (one “Yes, definitely”, nine “Yes, probably”; see Q.3a and b in Table 1). Only one of these had already decided to cultivate. A separate univariate BART model showed that willingness rose markedly when NVEI exceeded 17 (Figure 2b), with a PRAUC of 0.83 (Figure 3b).

These findings suggest that NVEI is a strong predictor of both actual decisions and willingness, supporting its use as a modelling tool for understanding farmer motivations.

### 3.2 Net expected impacts on specific goals driving decision and willingness

Posterior probability analysis identified which goal-specific NVEI components were most strongly correlated with either having made or being willing to make the cultivation decision (Figure 4).

**Figure 4.**
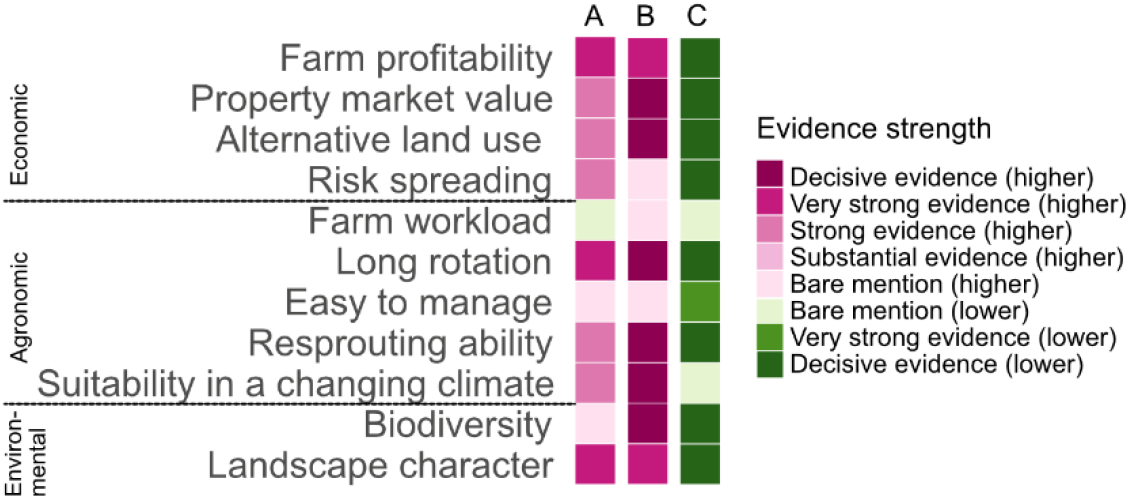
Posterior probabilities of differences in partial NVEI between respondents who had (A) decided or (B) were willing to cultivate *Populus* spp. on agricultural land and those who had not. Column (C) shows posterior probabilities for respondents who had explicitly answered “No, definitely not” willing to cultivate, compared with all other responses. For C, calculations were made excluding three missing values.

For actual decisions, very strong evidence supported the importance of net expected impacts on *Farm profitability* (economic), *Long rotation* (agronomic), and *Landscape character* (environmental). Other significant contributors included *Property market value, Alternative land use, Risk spreading, Resprouting ability, and Suitability in a changing climate*.

For willingness, decisive evidence supported the importance of net expected impacts on *Property market value, Alternative land use, Long rotation, Resprouting ability, Suitability in a changing climate*, and *Biodiversity*, with strong evidence for *Farm profitability* and *Landscape character*.

For those 29 respondents (28%) explicitly unwilling (“No, definitely not”), partial NVEI scores were decisively lower across most goals, except for *Farm workload* and *Suitability in a changing climate* (Figure 4C). Very strong evidence for lower scores was also found for the goal of *Easy to manage*.

Notably, *Farm workload* and *Easy to manage* scored low across nearly all respondent groups, including those willing to cultivate.

Pairwise comparisons (Figure 5) indicated that, among decision-makers, expected impacts of *Suitability in a changing climate* was prioritised over *Property market value* and *Farm workload*. For the willing group, expected impacts of *Alternative land use* and *Suitability in a changing climate* were more influential than *Profitability, Risk spreading*, and *Easy to manage*.

**Figure 5.**
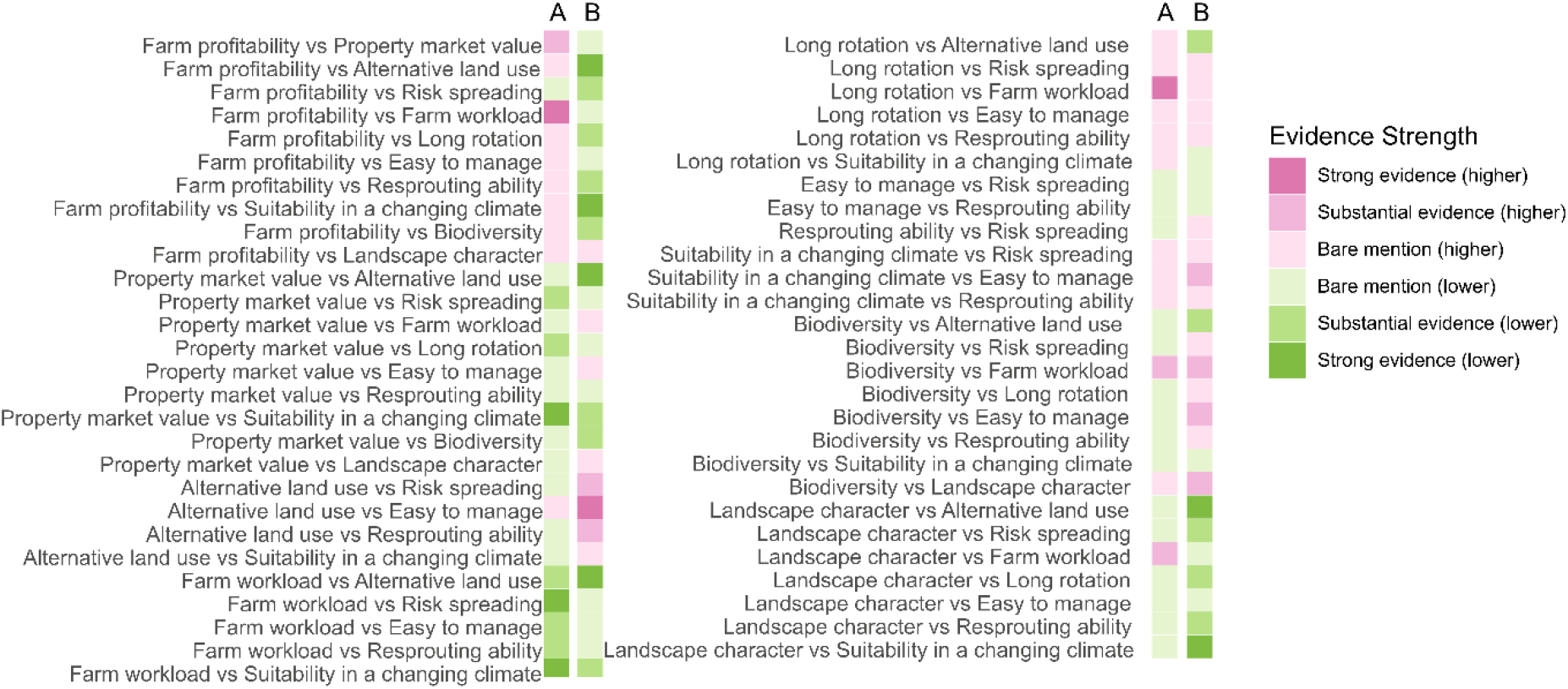
Posterior probabilities of differences in partial NVEI between each pair of goals for respondents who have (A) decided to cultivate or expand the cultivation of *Populus* spp. on agricultural land, and (B) are willing to do so. “Higher” indicates that goal 1 > goal 2, while “Lower” indicates that goal 2 > goal 1.

### 3.3. Hypothesis testing

Building on the model validation, we tested the empirical consequences of four hypotheses relating to differences in net values of expected impacts across goals (partial NVEI) and silvicultural experience.

#### 3.3.1 H.1

Net expected economic, agronomic, and environmental impacts and consequences of cultivating *Populus* spp. are correlated with actual decisions to cultivate or expand the cultivation of *Populus* spp. on agricultural land

##### H.1.a. Net expected economic consequences are positively correlated with the decision to cultivate <u>Populus</u> spp. (<u>cf</u>. Thomson Ek et al. 2024)

Strong to decisive evidence showed that expected values on economic goals, such as *Farm profitability*, are more strongly correlated with decisions to cultivate *Populus* spp. than with indecision or choosing not to cultivate them (Figure 4), and these goals had significantly higher partial NVEI in the Forested region compared to the Intermediate and Lowland regions (Figure 6). We found strong evidence that the expected consequences of *Populus* spp. cultivation on the goal of the *Property market value* were lower in the Lowland region than in the Intermediate region.

**Figure 6.**
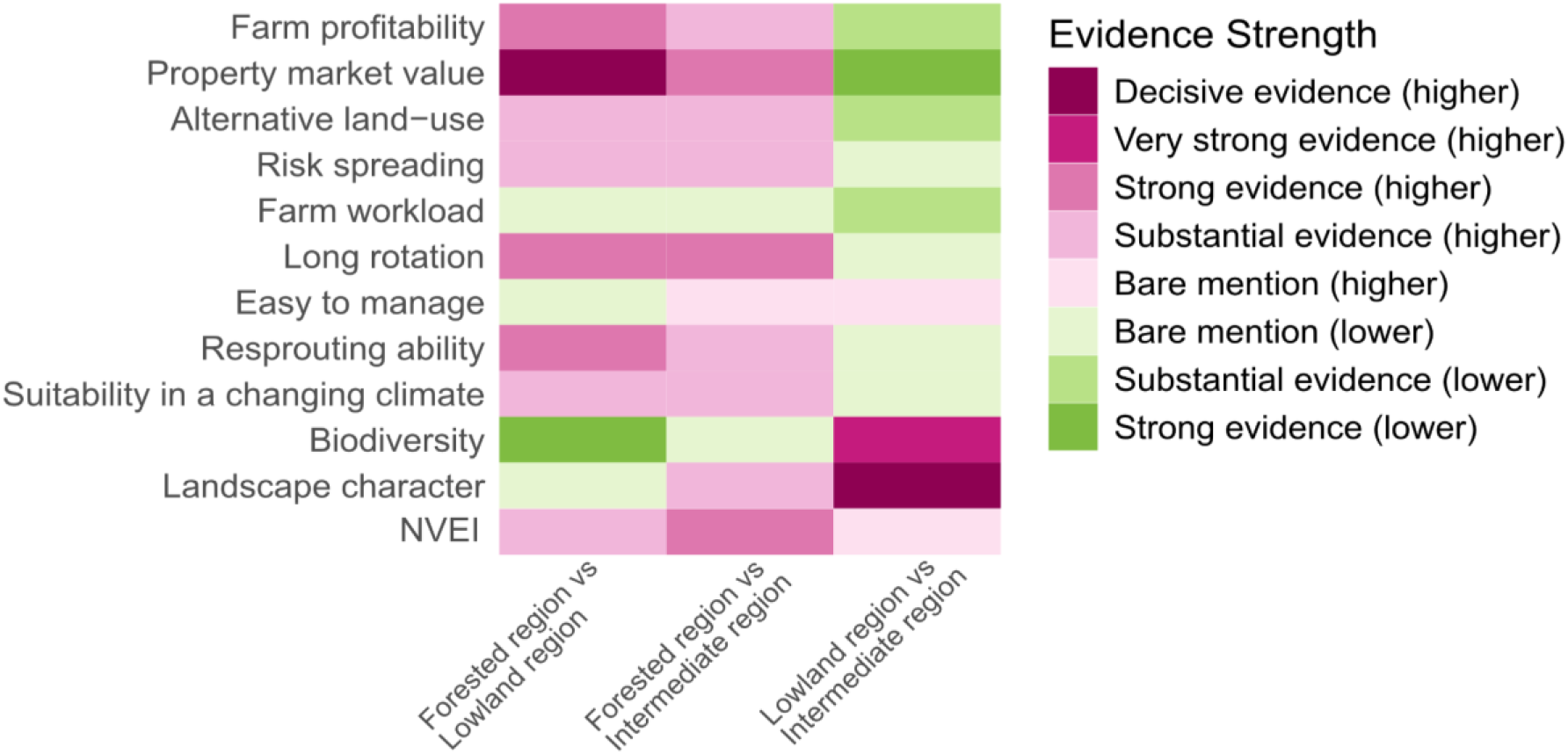
Posterior probabilities of differences in partial NVEI for each goal and the total NVEI between pairs of regions. The heatmap shows the probability that group A (one region) has a larger posterior probability than group B (another region).

Supporting this, rankings of expected values revealed that *Farm profitability* and *Risk spreading* were significantly more important contributors to NVEI than *Farm workload*, while *Suitability in a changing climate* was also ranked above *Property market value* (Figure 5).

##### H1b: Net expected agronomic impacts are negatively correlated with the decision to cultivate <u>Populus</u> spp. (<u>cf</u>. Paulrud and Laitila 2010)

We found very strong evidence supporting a positive correlation between net expected impacts on the agronomic goals of *Long rotation, Resprouting ability*, and *Suitability in a changing climate* and decision-makers, but not *Farm workload* and *Easy to manage* (Figure 4). *Long rotation* and *Resprouting ability* had higher partial NVEI in the Forested region compared to the Lowland (and for *Long rotation*, also compared to Intermediate) region (Figure 6).

##### H1c: Net expected environmental impacts are negatively correlated with the decision to cultivate <u>Populus</u> spp. (<u>cf</u>. Paulrud and Laitila 2010)

Net expected impacts on *Landscape character* were found to be strongly positively correlated with the decision to cultivate (Figure 4), while *Biodiversity* expectations were found to be significantly negatively correlated in the Forested region and positively correlated in the Lowland region (Figure 6). Conversely, *Landscape character* and *Biodiversity* expectations were decisively higher in the Lowland region compared to the Intermediate.

Although not top-ranked among decision drivers, *Landscape character* still contributed more to NVEI than *Farm workload*, highlighting its relative importance within environmental considerations (Figure 5).

#### 3.3.2 H2

Net expected economic, agronomic, and environmental impacts and consequences of cultivating *Populus* spp. are correlated with farmers’ willingness to cultivate *Populus* spp. on agricultural land

##### H2a: Net expected economic consequences are positively correlated with the willingness to cultivate <u>Populus</u> spp. (<u>cf</u>. Thomson Ek et al. 2024)

Decisive or very strong evidence indicated that expectations for most economic goals (excluding *Risk spreading*) were more positively evaluated by those willing to cultivate (Figure 4). These goals also had significantly higher NVEI scores in the Forested region, particularly for *Property market value* and *Farm profitability* (Figure 6).

In terms of relative importance, *Alternative land use* stood out. It was found to be significantly more strongly correlated with higher NVEI than *Farm profitability, Property market value, Farm workload, Easy to manage*, and *Landscape character* (Figure 5B).

##### H2b: Net expected agronomic impacts are negatively correlated with the willingness to cultivate <u>Populus</u> spp. (<u>cf</u>. Paulrud and Laitila 2010)

Respondents who were willing to cultivate expected decisively higher positive consequences of *Long rotation, Resprouting ability*, and *Suitability in a changing climate*, but not on *Farm workload* and *Easy to manage* (Figure 4). Regional analysis showed *Long rotation* and *Resprouting ability* expectations were higher in the Forested region compared to others, and *Farm workload* was lower in the Lowland region compared to the Intermediate region (Figure 6).

In pairwise comparisons, expected impacts of *Suitability in a changing climate* were rated significantly higher than *Farm profitability, Alternative land use*, and *Landscape character* (Figure 5B).

##### H2c: Net expected environmental impacts are negatively correlated with the willingness to cultivate <u>Populus</u> spp. on agricultural land (<u>cf</u>. Paulrud and Laitila 2010)

The partial NVEI of both *Biodiversity* and *Landscape character* were rated higher among those willing to cultivate (Figure 4), but *Biodiversity* expectations were lower in the Forested region compared to the Lowland region (Figure 6).

Although impacts on these environmental goals were positively valued overall, pairwise comparisons suggested that impacts of *Alternative land use* and *Suitability in a changing climate* were more influential contributors to NVEI than *Landscape character* (Figure 5B).

### 3.4. H.3

Previous experience with silviculture or tree cultivation is positively correlated with actual decisions to cultivate or expand the cultivation of *Populus* spp. on agricultural land

Bayesian proportions tests indicated strong evidence that forest landowners were more likely to have decided to cultivate *Populus* spp. than non-forest landowners (Table 3).

**Table 3.**
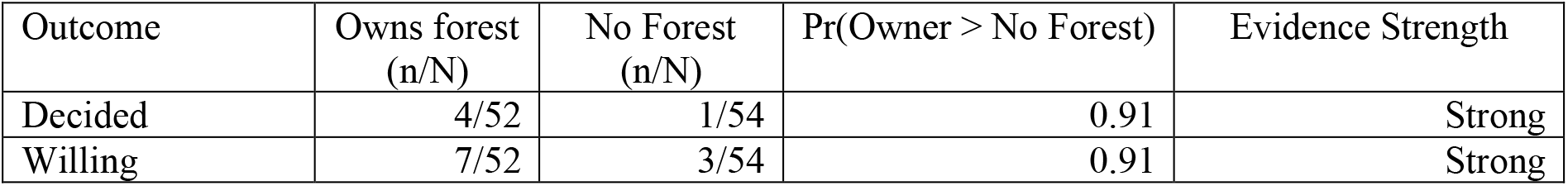
Bayesian proportions test results on the correlation between owning *Populus* spp. land and the decision or willingness to cultivate *Populus* spp.

| Outcome | Owns forest<br>(n/N) | No Forest<br>(n/N) | Pr(Owner > No Forest) | Evidence Strength |
| --- | --- | --- | --- | --- |
| Decided | 4/52 | 1/54 | 0.91 | Strong |
| Willing | 7/52 | 3/54 | 0.91 | Strong |

### 3.5. H.4

Previous experience with silviculture or tree cultivation is positively correlated with farmers’ willingness to cultivate *Populus* spp. on agricultural land

#### H4a Forest ownership is correlated with the willingness to cultivate or expand the cultivation of <u>Populus</u> spp. on agricultural land

We found strong evidence that forest landowners were more willing to cultivate *Populus* spp. in the coming decade than other landowners (Table 3).

#### H4b <u>Populus</u> spp. land ownership is correlated with the willingness to cultivate <u>Populus</u> spp. on agricultural land

Very strong evidence supported that ownership of existing *Populus* plantations was correlated with willingness to cultivate *Populus* spp. on agricultural land compared to non-owners (Table 4).

**Table 4.** Results of a Bayesian proportions test testing whether owning *Populus* spp. land is positively correlated with being willing to cultivate *Populus* spp. in the coming decade.

| Outcome | Owns *Populus* spp.<br>(n/N) | No *Populus* spp.<br>(n/N) | Pr(Owner > No<br>*Populus* spp.) | Evidence Strength |
| --- | --- | --- | --- | --- |
| Willing* | 3/12 | 7/90 | 0.97 | Very strong |

\* One respondent answering *Don't know* was excluded from the test.

#### 3.6. Communication needs of respondents regarding the cultivation of *Populus* spp. on agricultural land

Respondents in different regions evaluated the impacts of *Populus* spp. cultivation differently, reflecting distinct informational needs (Figure 6). In the Lowland region, higher partial NVEI scores for environmental goals, particularly *Landscape character* and *Biodiversity*, indicate that farmers expect *Populus* spp. to contribute positively to these outcomes. Communication efforts in this region can build on this expectation by providing more specific, practical information on how *Populus* spp. cultivation supports habitat diversity, visual landscape values, and ecological connectivity within farming systems.

In contrast, respondents in the Forested region rated economic and agronomic expectations, such as *Farm profitability, Long rotation*, and *Resprouting ability*, more positively. Environmental expectations like *Biodiversity* received notably lower NVEI scores in this region, not necessarily because they are considered unimportant, but likely due to lower expectations that *Populus* spp. supports these outcomes. This suggests a knowledge gap rather than a lack of concern. Here, communication could provide comparative evidence, for example, showing how *Populus* spp. stands differ from spruce monocultures in terms of understory diversity or ecosystem function. Clarifying such differences may help farmers assess the environmental implications of species choice more accurately.

Notably, *Farm workload* and *Easy to manage* received low scores across nearly all respondent groups, including those willing to cultivate. These low expectations suggest that practical aspects of labour and management may not be well understood, are considered irrelevant, or are underestimated in terms of their relevance. Expectations on workload and management ease may be more amenable to clarification through experience, peer exchange, or targeted extension materials.

A substantial minority of respondents (28%) explicitly indicated that they would “definitely not” consider *Populus* spp. cultivation, and this group showed consistently low partial NVEI scores across almost all goal components. While this pattern suggests limited openness to change, it still points to the importance of making balanced, transparent, and evidence-informed information available, without assuming persuasion is possible or appropriate.

Overall, these findings indicate that effective communication about *Populus* spp. cultivation should be regionally tailored, not to shift values, but to address specific knowledge needs. In the Lowland region, this means deepening understanding of the environmental functions of *Populus* spp.; in the Forested region, it involves clarifying economic and agronomic benefits while addressing gaps in understanding about its environmental impacts, especially relative to species like spruce. Across all regions, clear guidance on management practices and system integration can support more informed decision-making among farmers.

## 4. Discussion

We developed a methodology to identify key factors shaping farmer behaviour, with a particular focus on awareness and motivation gaps that may hinder broader adoption of *Populus* spp. cultivation on agricultural land for biomass energy production. By analysing decision-making processes and willingness to cultivate *Populus* spp., our models based on Net Value of Expected Impact (NVEI) successfully captured critical dimensions of farmers’ reasoning (Figure 2).

The emerging patterns suggest that NVEI is a useful framework for understanding cultivation decisions, even though we did not conduct a formal hypothesis test of predictive performance. Nevertheless, the models demonstrated good predictive accuracy, as indicated by PRAUC values (Figure 3), supporting their relevance in explaining observed behaviours.

Our approach builds on models like the agent-based framework by Pralhad et al. (2021) by directly integrating empirical data on farmers’ expected utilities and decision-making criteria. Unlike simulation models that rely on assumptions about preferences, we quantify farmers’ expected utilities across economic, agronomic, and environmental dimensions, revealing regional and experiential differences. This empirical basis enables more precise identification of communication needs and tailored strategies to support communications, addressing a gap in models that focus primarily on adoption rates rather than underlying motivations.

As in many studies involving farmer decision-making and land-use preferences (e.g., Rommel et al. 2022), our survey encountered a relatively low response rate. While this limits the statistical generalisability of the findings, prior literature suggests that low participation is a common feature of studies focused on novel or long-term land-use changes. Moreover, the presence of distinct patterns across regions and decision types suggests that the data still provides valuable insights into farmer motivations and potential barriers.

We acknowledge a potential interdependence between the auxiliary hypothesis (establishing the predictive power of NVEI on behaviour through BART models, see Figure 3) and the main hypotheses (H.1 and H.2). By first confirming NVEI’s predictive relevance, we accounted for this interrelationship when interpreting the main hypothesis results, thereby integrating the dependency into our overall conclusions.

The following hypotheses and their empirical consequences were used to structure the analysis and interpretation:

### 4.1. H.1

Net expected economic, agronomic, and environmental impacts and consequences of cultivating *Populus* spp. are correlated with actual decisions to cultivate or expand the cultivation of *Populus* spp. on agricultural land.

We tested the empirical consequences of hypothesis H.1, which states that net expected economic, agronomic, and environmental impacts and consequences of cultivating *Populus* spp. are correlated with actual decisions to cultivate or expand the cultivation of *Populus* spp. on agricultural land. To test the hypothesis, we used a structured set of expected outcomes within each of the three categories: economic, agronomic, and environmental.

The test of H.1.a, stating that net expected economic consequences are positively correlated with the decision to cultivate *Populus* spp., and the test of H.1.b, stating that net expected agronomic impacts are negatively correlated with the decision to cultivate *Populus* spp., yielded strong to decisive evidence in support of H.1.a but not for H.1.b. Consequences related to farm profitability and property market value (economic), but also agronomic considerations such as *Long rotation* periods and *Resprouting ability*, were particularly influential in shaping actual cultivation decisions. However, we did not find any strong correlation with the agronomic goals of *Farm workload* and *Easy to manage*, which consistently received low expected impact scores, even among those who had already decided to cultivate.

In the Forested region, where the fertility of the soil and price of land are lower (Swedish Board of Agriculture 2019), economic and certain agronomic factors appeared especially salient, suggesting that farmers expect profit and agronomic suitability from *Populus* spp. cultivation. These findings align with earlier research showing that farmer adoption of biomass crops is influenced not only by biophysical constraints but also by perceived economic advantages (Schultze et al. 2016). They also support Thomson Ek’s (2024) observation that clear economic benefits are crucial in contexts of price sensitivity and uncertain markets. That said, the consistently low evaluation of labour-related agronomic factors points to an important communication need: farmers may lack adequate information about the practical implications of *Populus* spp. cultivation. Addressing knowledge gaps on workload and management could further support adoption, especially among undecided farmers.

The test of H.1.c, stating that net expected environmental impacts are negatively correlated with the decision to cultivate *Populus* spp. was also corroborated. However, the relationship between expected impacts on environmental goals and cultivation decisions varied by region. In the Forested region, expected impacts on environmental goals, such as biodiversity, were indeed negatively correlated with actual cultivation decisions, whereas in the Lowland region, expected impacts on these goals were found to be positively correlated. This suggests that while expected impacts on environmental goals can act as a barrier in some contexts, they may serve as a motivating factor in others. A study by Lindbladh et al. (2014) compared young hybrid aspen stands with spruce plantations of similar age in southern Sweden. The findings revealed that hybrid aspen stands supported higher bird species richness and abundance, as well as a distinct community composition. This increased biodiversity is attributed to the greater structural complexity and tree species diversity in aspen stands, which provide more varied habitats for birds. For understory vegetation in *Populus* spp. plantations on former agricultural land in southern and central Sweden, Weih et al. (2003) found that species richness in these plantations was comparable to that of old-growth mixed deciduous forests in central Sweden, although slightly lower than in southeastern regions. This suggests that *Populus* spp. plantations can contribute positively to plant biodiversity, especially when established on previously intensively managed agricultural lands.

Because only H.1.a and H.1.c were corroborated, while H.1.b was not, H.1 was partly corroborated. This indicates that while expectations on economic and some environmental factors significantly influence cultivation decisions, expected impacts on agronomic factors do not consistently exert a negative influence as hypothesised. Thus, although previous studies (e.g. Paulrud and Laitila 2010, Hand and Tyndall 2018) emphasise concerns about the agronomic suitability of energy crops, our findings show a more complex pattern, with certain agronomic traits (e.g. *Long rotation, Resprouting ability*) seen as advantages by some decision-makers. This partly aligns with earlier observations by Hand and Tyndall (2018), who found that many farmers in the U.S. Great Plains held sceptical or negative views toward the cultivation of trees for bioenergy. In our study, a similar scepticism appeared in the Forested region, likely contributing to the lack of support for H.1.b. In contrast, in the Lowland region, the expected environmental benefits of *Populus* spp. cultivation may have outweighed concerns, contributing to more favourable decisions.

### 4.2 H.2

Net expected economic, agronomic, and environmental impacts and consequences of cultivating *Populus* spp. are correlated with farmers’ willingness to cultivate *Populus* spp. on agricultural land

The test of the empirical consequence H.2.a, stating that net expected economic consequences are positively correlated with willingness to cultivate *Populus* spp. on agricultural land, found decisive to very strong support for several economic goals, such as *Property market value* and *Farm profitability*. While economic motivations such as profitability and property value were found to be strongly correlated with willingness, the expected consequences on *Risk spreading*, often assumed to be a motivation for diversifying into biomass crops, did not contribute to willingness in our sample. This finding aligns with earlier work suggesting that the adoption of bioenergy crops involves complex risk considerations that may outweigh the perceived benefits of diversification (Skevas et al. 2016). Rather than functioning as a straightforward strategy to mitigate production risks, diversification into energy crops may introduce new forms of uncertainty, particularly when market structures are underdeveloped or agronomic knowledge is limited.

Regarding H.2.b, stating that net expected agronomic consequences are negatively correlated with the willingness to cultivate *Populus* spp., the evidence was mixed. While expectations on *Long rotation, Resprouting ability*, and *Suitability in a changing climate* were positively correlated with willingness to cultivate, we found no support for the agronomic factors of *Farm workload* and *Easy to manage*. Thus, H.2.b was not corroborated, and H.2 was partially corroborated. These results suggest that while some agronomic goals are seen as advantages, others may not be fully understood or valued, possibly due to limited operational knowledge. Communication strategies targeting prospective cultivators could benefit from addressing these specific knowledge gaps.

The test of the empirical consequence H.2.c, stating that net expected environmental impacts are negatively correlated with willingness to cultivate *Populus* spp., also did not find support. In fact, in the Lowland region, we observed a positive correlation between willingness and the expected impacts on the environmental goals of *Biodiversity* and *Landscape Character*. This directly contradicts the expectation of H.2.c and further indicates that H.2 was only partially corroborated.

As discussed in H.1.c, ecological studies (e.g., Lindbladh et al. 2014; Weih et al. 2003) show that *Populus* plantations can provide greater biodiversity benefits compared to spruce monocultures common in the Forested region. The negative correlation between expected impacts on environmental goals and willingness observed in the Forested region may therefore reflect a lack of knowledge among farmers of these comparative ecological advantages. Given that spruce dominates the landscape there, farmers may not expect *Populus* spp. to be an environmentally beneficial alternative.

Addressing this information gap through targeted communication highlighting the superior biodiversity value of *Populus* spp. relative to spruce could help increase willingness to cultivate in the Forested region.

These findings diverge from earlier studies. For example, Paulrud and Laitila (2010) highlighted that non-economic factors, especially scepticism toward taller, long-rotation crops, reduced expected utility among Swedish farmers. Similarly, Hand and Tyndall (2018) documented reluctance among U.S. Great Plains farmers to convert farmland to bioenergy trees. However, our results suggest that such scepticism is not universal. In the Lowland region, in particular, expected impacts on environmental goals appear to support rather than hinder willingness. This underscores the importance of regional variation in farmers’ expectations of land use and cautions against overgeneralising farmer reluctance toward *Populus* spp. cultivation.

### 4.3 H.3

Previous experience with silviculture or tree cultivation is positively correlated with actual decisions to cultivate or expand the cultivation of *Populus* spp. on agricultural land

The test of H.3, stating that previous experience with silviculture or tree cultivation is positively correlated with actual decisions to cultivate or expand the cultivation of *Populus* spp. on agricultural land, found substantial evidence that forest ownership or prior *Populus* cultivation is positively correlated with actual decisions to cultivate. H.3 was therefore corroborated.

### 4.4 H.4

Previous experience with silviculture or tree cultivation is positively correlated with farmers’ willingness to cultivate *Populus* spp. on agricultural land

Tests of the empirical consequence H.4.a, stating that forest ownership is correlated with willingness to cultivate or expand the cultivation of *Populus* spp. on agricultural land, and H.4.b, stating that *Populus* spp. land ownership is correlated with the willingness to cultivate or expand the cultivation of *Populus* spp. on agricultural land, showed strong positive correlation between forest ownership and willingness, and also a strong correlation for those already owning *Populus* land. Therefore, H.4 was corroborated.

### 4.5 Implications for distinguishing between willingness and decisions

One key finding of this study is the divergence in motivations between farmers who have made decisions and those who are willing to do so. While willingness was influenced by expected impacts on a broad range of goals, including environmental ones, actual decisions were more narrowly aligned with economic and agronomic considerations. This underscores the importance of analysing not only willingness but, importantly, actual decisions, rather than treating the former as a simple proxy for the latter.

It is, however, important to treat self-reported measures such as the questionnaire responses used in this paper with caution, as participants may report values that align with their responses to present a coherent picture of their decision process. The direction of causality is therefore not certain: the reported expectations may have generated reported decisions, but these decisions may also have shifted participants’ expectations. Such shifts are particularly likely to appear for participants who have decided to plant *Populus* spp., and for those who report that they will not plant *Populus* spp. (*<u>cf</u>*. Ariely and Norton 2007).

### 4.6 Policy implications

Our findings have several implications for policymakers, advisory bodies, and extension services seeking to facilitate informed decisions about *Populus* spp. cultivation on agricultural land. These guidelines can inform the design of communication strategies, extension programs, and potential policy instruments, while aligning with principles of transparency and respect for farmers’ autonomy.

#### 4.6.1 Guidelines for effective communication

Based on our findings, we propose the following communication guidelines to support informed and autonomous decision-making about *Populus* spp. cultivation. Communication efforts should prioritise addressing knowledge gaps, especially regarding ecological and operational outcomes, rather than relying on value-based messaging. We emphasise that *Populus* spp. cultivation is not universally suitable and may, in some contexts, conflict with biodiversity conservation or food production priorities. The aim is not to promote *Populus* spp., but to provide accessible, balanced information to support land-use decisions.

##### Segment the audience by region and decision status

In the Forested region, where expectations on economic and certain agronomic goals (e.g. profitability, rotation length, resprouting ability) were prioritised, farmers may lack detailed knowledge of how *Populus* spp. compares to familiar species such as spruce. Communication should therefore focus on closing potential knowledge gaps, for example, by providing practical information on growth performance, management demands, and resilience under local conditions.

Expected impacts on ecological outcomes were also less emphasised in this region, likely due to the dominance of spruce monocultures and limited awareness of species-specific biodiversity impacts. Comparative data from studies such as Lindbladh et al. (2014) and Weih et al. (2003) could help clarify how *Populus* spp. plantations differ ecologically from more commonly used species.

In the Lowland region, farmers expected more positive impacts on environmental and aesthetic values. Here, communication can support interest in *Populus* spp. by providing information on how such plantations may contribute to biodiversity, habitat connectivity, and landscape character, but should also address practical considerations that may not yet be well understood.

For farmers who are already willing, messaging should focus on closing the gap between intention and action by addressing practical barriers such as access to planting material, management advice, and market information.

Among farmers who were firmly opposed, our results showed generally low expected impacts of *Populus* spp. across most goals. In these cases, communication should avoid persuasion and instead offer neutral, comparative information on land-use options. Respecting these views while ensuring access to reliable information helps maintain trust, even where interest is unlikely to change.

##### Prioritise closing knowledge gaps

Several decision goals, particularly *Farm workload* and *Easy to manage*, received consistently low expected values across all groups. Rather than reflecting a lack of interest, this may indicate limited knowledge of what *Populus* spp. cultivation entails in practice. Communication should aim to clarify these aspects by providing practical, experience-based information about labour intensity, machinery needs, seasonal tasks, and potential risks.

Similarly, the lower expectations on environmental goals in the Forested region may not reflect disinterest but rather a lack of species-specific ecological information. Communication can help bridge this gap by providing comparisons between *Populus* spp. and dominant species like spruce in terms of biodiversity and ecosystem service outcomes.

In contrast to more firmly held values (e.g. profitability, landscape preferences), expectations around management demands or ecological benefits may change when decision-makers have access to credible, targeted information. Tailored extension materials, peer learning opportunities, and site visits can help address these specific knowledge needs.

##### Highlight potential conflicts and trade-offs

In regions with highly productive cropland, allocating land to *Populus* spp. may involve trade-offs with food production. These should be communicated clearly, using relevant examples to show how others have evaluated and managed such trade-offs. This supports realistic expectations and better-aligned decision-making, without advocating for a specific land use.

Where biodiversity benefits are expected, particularly in the Lowland region, communication should clarify how these benefits vary with site, management, and scale. In the Forested region, contextualising biodiversity comparisons between *Populus* and existing spruce plantations may be especially important.

##### Leverage experiential knowledge

Farmers with forestry or *Populus* spp. cultivation experience reported higher levels of willingness and positive expectations. To support others, communication should include peer-to-peer learning, field visits, and testimonials.

##### Support informed choice with transparent, site-relevant data

Where interest exists, communication should offer regionally relevant information on *Populus* spp. performance, such as how it responds to different soil types, land-use histories, and climatic conditions. While some aspects of future performance may remain uncertain, clearly distinguishing between knowns and unknowns is essential. This supports farmer autonomy and avoids creating a sense of guided or predetermined outcomes.

By addressing knowledge gaps, clarifying operational aspects, and offering balanced comparisons, communication can empower farmers to make informed land-use decisions that align with their goals, contexts, and constraints, without attempting to persuade or steer them toward *Populus* spp. cultivation.

## 5. Conclusion

This study highlights the complexity of decision-making around *Populus* spp. cultivation, shaped by a mix of economic expectations, environmental values, and contextual realities. While economic concerns consistently influence decisions, the role of environmental and agronomic factors is more variable and often under-recognised. Importantly, the findings show that willingness to cultivate does not necessarily lead to action, pointing to the critical need for better alignment between available information and the practical needs of farmers.

Beyond its empirical contributions, the study demonstrates the value of decision-analytic methods and frameworks, particularly partial NVEI and Bayesian modelling, for uncovering how farmers weigh competing goals and uncertainties. These tools not only reveal where assumptions fail but also help identify where knowledge gaps are most likely to shape or stall decisions.

A key takeaway is that improved communication is not just about raising awareness but about enabling evaluation. Farmers need access to clear, regionally relevant, and experience-informed knowledge that addresses specific concerns such as biodiversity impacts, ease of management, and effects on workload. Current knowledge and outreach efforts often overlook these practical dimensions, limiting farmers’ ability to assess fit within their contexts.

Looking ahead, communication strategies should focus on building usable knowledge: credible content, contextualised, and directly applicable to farm-level decisions. Rather than uniformly promoting adoption, communication should support farmers in making informed, autonomous choices based on the realities of their land, goals, and constraints.

For researchers and policymakers alike, this means shifting focus from influencing attitudes to enabling decisions. By prioritising the co-development and targeted delivery of actionable knowledge, future efforts can more effectively support the voluntary and context-sensitive adoption of sustainable land-use practices.

## Acknowledgements

We sincerely appreciate all the interviewees and questionnaire participants in Scania County for their invaluable input. We also extend our thanks to the Swedish Farmers Association for their assistance in enabling the participation of potential respondents and to the Swedish Board of Agriculture for providing access to relevant data.

## Declaration of generative AI and AI-assisted technologies in the writing process

During the preparation of this work, the authors used ChatGPT in order to edit the English language. After using this tool/service, the authors reviewed and edited the content as needed and take full responsibility for the content of the published article.

## Data availability

The data can be accessed from the Swedish National Data Repository () by anyone with a legitimate interest in the data, as long as the transfer of data complies with the Swedish and European regulations on data protection.

